# Cytoarchitectonic similarity is a wiring principle of the human connectome

**DOI:** 10.1101/068254

**Authors:** Alexandros Goulas, René Werner, Sarah F Beul, Dennis Säring, Martijn van den Heuvel, Lazaros C Triarhou, Claus C Hilgetag

**Affiliations:** Department of Computational Neuroscience, University Medical Center Hamburg-Eppendorf, Martinistr. 52, 20246 Hamburg, Germany; Brain Center Rudolf Magnus, Department of Psychiatry, University Medical Center Utrecht; Laboratory of Theoretical and Applied Neuroscience, University of Macedonia, Egnatia 156, Thessaloniki 54636, Greece; Department of Health Sciences, Boston University, 635 Commonwealth Ave., Boston, MA 02215, USA

## Abstract

Understanding the wiring diagram of the human cerebral cortex is a fundamental challenge in neuroscience. Elemental aspects of its organization remain elusive. Here we examine which structural traits of cortical regions, particularly their cytoarchitecture and thickness, relate to the existence and strength of inter-regional connections. We use the architecture data from the classic work of von Economo and Koskinas and state-of-the-art diffusion-based connectivity data from the Human Connectome Project. Our results reveal a prominent role of the cytoarchitectonic similarity of supragranular layers for predicting the existence and strength of connections. In contrast, cortical thickness similarity was not related to the existence or strength of connections. These results are in line with findings for non-human mammalian cerebral cortices, suggesting overarching wiring principles of the mammalian cerebral cortex. The results invite hypotheses about evolutionary conserved neurobiological mechanisms that give rise to the relation of cytoarchitecture and connectivity in the human cerebral cortex.

## Introduction

A major endeavor in neuroscience aims at mapping and understanding the macroscale connectivity of the human brain by using *in vivo* imaging techniques (e.g., Hagmann et al., 2008; Van Essen et al., 2013). This approach for resolving the so-called human connectome conceptualizes the brain as a collection of nodes, such as cortical divisions defined by macroscopic or microscopic criteria (e.g., morphological landmarks or cytoarchitectonic differentiation) and edges, such as diffusion MRI tractography streamlines linking the different divisions. Previous studies have uncovered key topological features of the human connectome, such as the presence of hierarchically organized modules (Hilgetag et al., 2000; Meunier et al., 2009) or highly connected hub nodes which are also tightly interconnected among themselves, forming so-called rich-clubs (van den Heuvel and Sporns, 2013; Bullmore and Sporns, 2009). The exact functional role of such topological features of the structural connectome continues to be elucidated through empirical and computational approaches (e.g., van den Heuvel and Sporns, 2013; Zamora-López et al., 2010; Senden et al., 2014). The significance of the properly configured connectome for healthy brain functioning is underlined by evidence suggesting that topological features such as hubs are affected in neurological disorders (e.g., Crossley et al., 2014). Despite such advances in mapping higher order topological properties of the human connectome, we still have an incomplete understanding of the basic principles of its organization. For example, it is unknown why certain cortical areas are interconnected while others are not, giving rise to the characteristic pattern of cortico-cortical connections of the human brain.

Useful insights originate from classic cytoarchitectonic and invasive connectivity studies in animal models, such as the macaque monkey. In a series of studies spanning several decades, Pandya, Barbas and colleagues have demonstrated that patterns of inter-areal connections and gradients of cytoarchitecture across the macaque cortex are related (Yeterian and Pandya, 1985; Pandya, Seltzer and Barbas, 1988; Pandya and Yeterian, 1990). More specifically, qualitative evidence suggests that areas with similar cytoarchitecture tend to be connected, indicated a “similar prefers similar” wiring principle of cortico-cortical connectivity (for recent reviews see Barbas, 2015; Pandya et al., 2015). Our previous work also demonstrated that such a cytoarchitecture-based principle can explain connectional features of the cerebral cortex, such as the existence, strength and laminar patterns of connections in several mammalian species, particularly the cat (Hilgetag and Grant, 2010; Beul et al., 2015a), mouse (Goulas et al., in press) and macaque (Barbas et al., 2005; Beul et al., 2015b; Hilgetag et al., 2016). Moreover, topological features such as the number of connections of an area also appear related to cytoarchitectonic features including neuronal density (Scholtens et al., 2014; Beul et al., 2015a; 2015b; van den Heuvel et al., 2015a). These findings suggest that a “similar prefers similar” wiring principle might operate across mammalian species. Here we explicitly examined if such a similarity principle also applies to fundamental features of the human connectome.

Information on the cytoarchitecture of the human cortex is obtained with high resolution by post-mortem histology, a field with a long tradition (e.g., Campbell, 1905; Brodmann, 1909; von Economo and Koskinas, 1925). In particular, the work of von Economo and Koskinas (1925) provided detailed measurements of various structural traits of human cortical areas, such as their layer thickness and cell density. Despite recent efforts in the cytoarchitectonic mapping of human cortical areas in stereotaxic space (e.g., Eickhoff et al., 2005), there exists as yet no comprehensive quantitative map of the cytoarchitecture of human cerebral cortical areas. Promising projects such as the BigBrain project (Amunts et al., 2013) are intended to fill this gap, but are still in their infancy. Therefore, the work of von Economo and Koskinas, while bound to the limitations and scientific practices of its era, is at this point still the only comprehensive source for quantitative cytoarchitectonic information on the human cerebral cortex.

Along with investigating cytoarchitectonic similarity, we aimed at examining whether the cortical thickness of regions was related to any of the basic features of the human connectome. Cortical thickness varies systematically across cortical areas and is related to the cytoarchitecture of cortical areas. Specifically, less differentiated areas of lower neuronal density are thicker, whereas more differentiated, denser areas are thinner (e.g., von Economo and Koskinas, 1925). Therefore,we examined conjointly if and how cytoarchitecture and thickness relate to the existence and strength of cortical connections, as in previous studies of the macaque cortex (Beul et al., 2015b; Hilgetag et al., 2016).

## Materials and Methods

### Constructing the human structural connectome

We constructed the human connectome from diffusion-weighted data of the Human Connectome Project (HCP) release Q3 (http://www.humanconnectome.org/documentation/Q3/) (Van Essen et al., 2013), which data constitutes a state-of-the-art connectivity set based on a large population of subjects. Specifically, this dataset includes 215 subjects with an age range of 22 to 35 years. Imaging parameters were: voxel size 1.25 mm isotropic, TR/TE 5520/89.5 ms, 270 diffusion directions with diffusion weighting 1000, 2000, or 3000 s/mm^2^. Diffusion weighted imaging data processing included eddy current and susceptibility distortion correction, reconstruction of the voxel-wise diffusion profile using generalized q-sampling imaging, and whole-brain streamline tractography. Cortical segmentation was performed based on the high-resolution T1-weighted image (voxel size: 0.7 mm isotropic) of each subject using FreeSurfer (Fischl et al., 2004). The cortical sheet of each subject was subsequently parcellated in 34 regions per hemisphere based on the Desikan-Killiany atlas (Desikan et al., 2006). These regions constituted the nodes for the connectome reconstruction for each subject. The strength of the connection between a pair of regions was taken as the number of streamlines linking the two regions. This procedure resulted in a 68x68 weighted and undirected matrix per subject (215 subject-specific connectomes in total). For the analysis of the existence of connections, they were treated in a binary fashion, that is, as present or absent, irrespective of their weight. For constructing a group average connectome, the connections that were present in at least 66% of the subjects were taken into consideration (Figure 1). In the absence of the ground truth for the whole brain connectional architecture of humans, this threshold was based on pragmatic recommendations for constructing group-average connectomes (de Reus and van den Heuvel, 2013). Using a different threshold (in the range of 60-75%) for constructing the group average connectome did not affect the results. Moreover, we performed a subject-wise analysis as described below.

**Figure 1.**
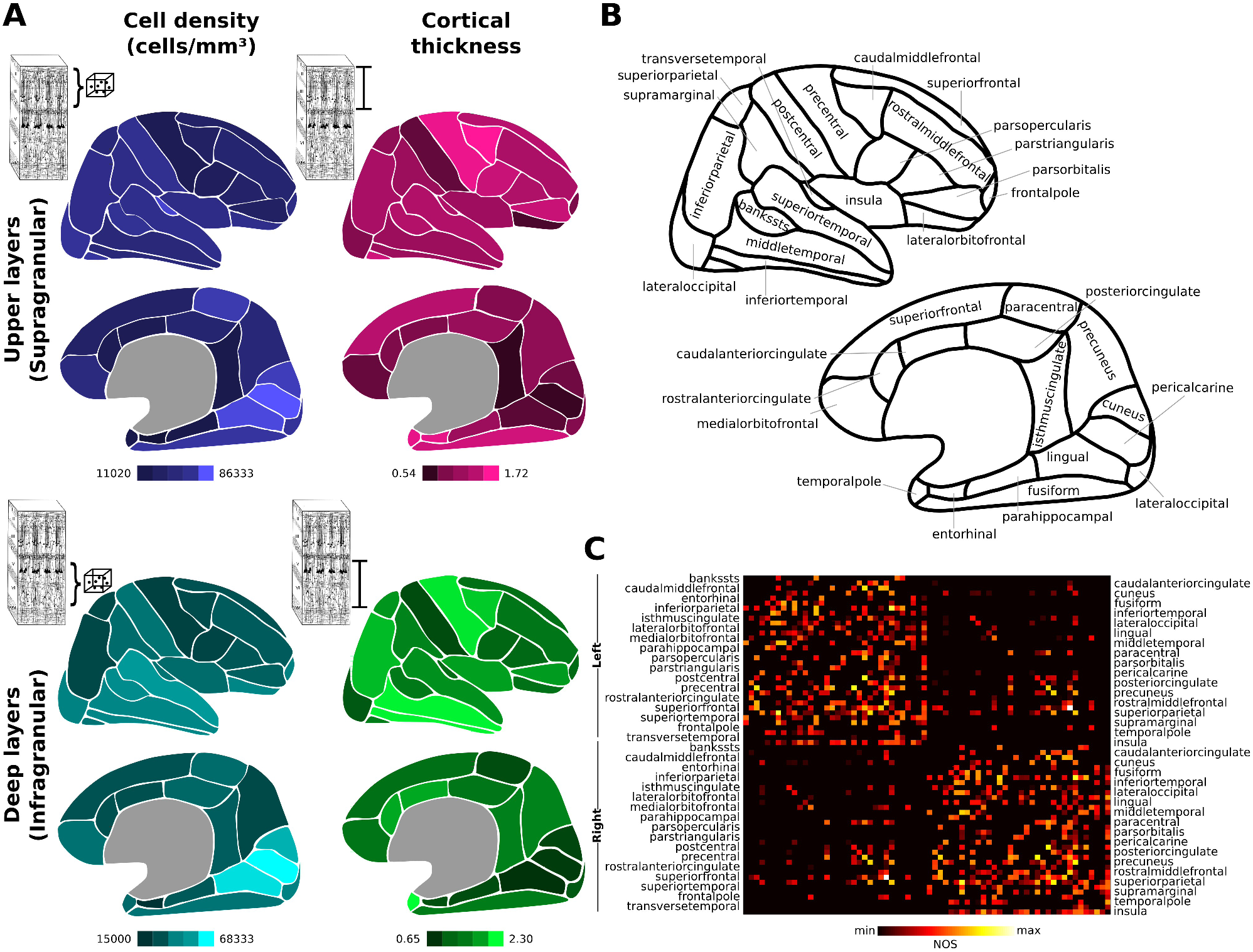
Regions of the Desikan-Killiany atlas, their structural traits and the connectome. **A**. Regions of the Desikan-Killiany atlas were matched based on topographic information and macroscopic landmarks to cortical areas of the von Econo and Koskinas atlas (see Table 1) and the respective infra- and upper cell density per mm^3^ and thickness was assigned to each region. **B.** Schematic depiction of the regions and names of the Desikan-Killiany atlas. **C.** Whole brain connectome assembled based on the Desikan-Killiany atlas and diffusion imaging data from the Human Connectome Project.

**Table 1.**
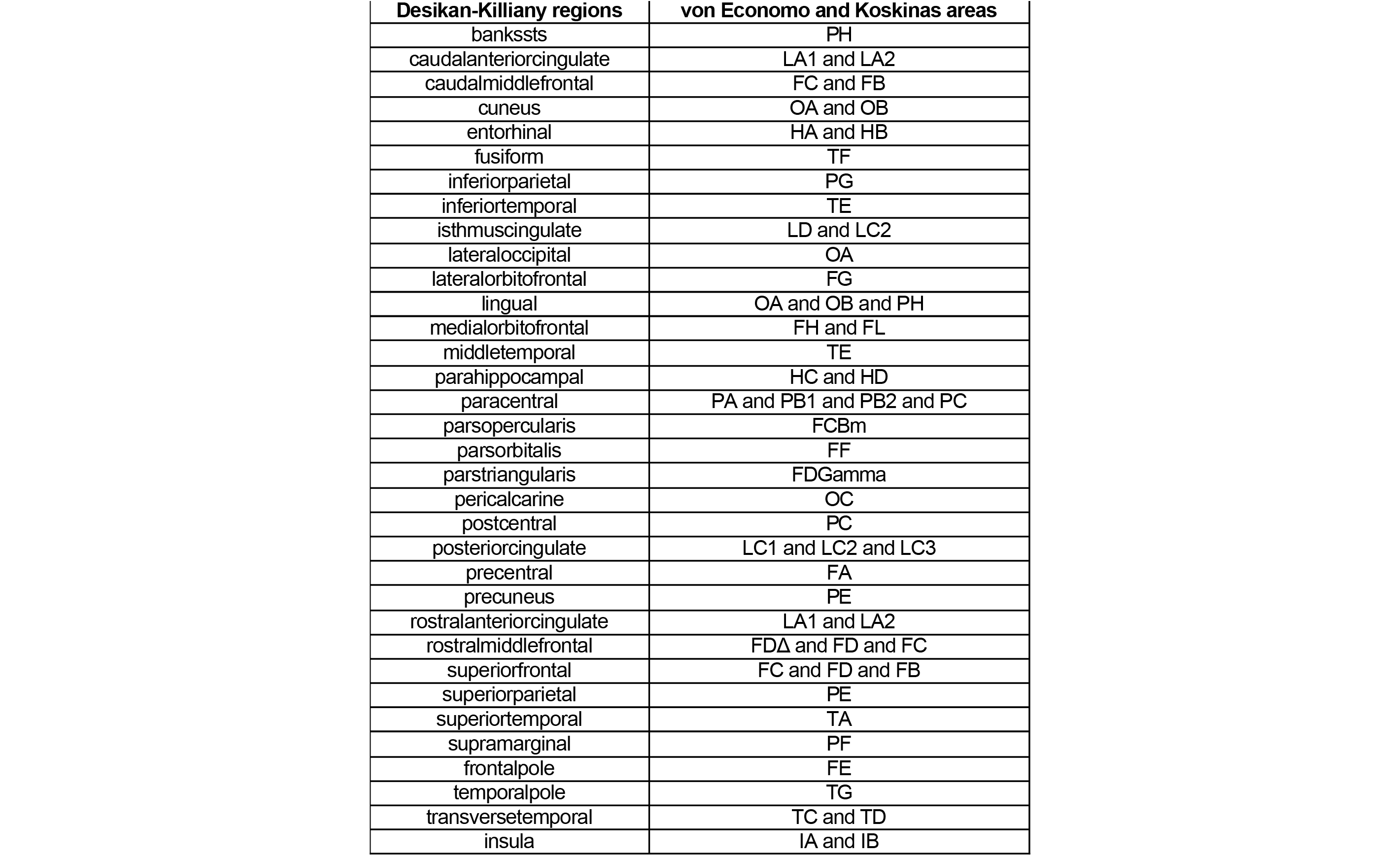
Correspondence of the Desikan-Killiany regions to von Economo and Koskinas “ground”, “variant” or “modification” areas. These matches were used to assign the structural traits (deep and upper cell density per mm^3^ and thickness) to the Desikan-Killiany regions.

### Matching the von Economo and Koskinas cortical areas to the macroscopic-based Desikan-Killiany atlas

We aimed at relating structural traits (laminar thickness and cytoarchitecture) to the presence or absence of connections between human cortical regions. We employed the cell density per mm^3^ as a quantitative measure of the characteristic cytoarchitecture of cortical regions (e.g., Dombrowski et al., 2001; Medalla and Barbas, 2006; Beul et al., 2015b). The only currently available, comprehensive source for this information for the human brain is the work of von Economo and Koskinas (1925). The von Economo and Koskinas atlas is only available as a schematic 2D drawing. Therefore, the first step of our approach involved the matching of the cortical areas of the von Economo and Koskinas atlas to the Desikan-Killiany atlas (Desikan et al., 2006), as in previous analyses (van den Heuvel et al., 2015a). The Desikan-Killiany atlas is available in stereotaxic space and can be used for assembling the human connectome (e.g., Hagmann et al., 2008; van den Heuvel et al., 2015a). This atlas contains 34 macroscopically defined regions for each hemisphere. The drawings of the von Economo and Koskinas atlas from von Economo (1927) and macroscopic landmarks were used as guides for matching the von Economo and Koskinas cortical areas to the Desikan-Killiany regions (see Table 1 for corresponding regions). The quantitative measurements tabulated in von Economo and Koskinas (1925) on the thickness and cell density per mm3 on the laminar compartments of the von Economo and Koskinas cortical areas were then used to construct deep and upper layer profiles for the regions of the Desikan-Killiany atlas.

We constructed separate infragranular (deep layer) and supragranular (upper layer) region profiles, because cytoarchitectonic gradients, and thus cytoarchitectonic differences, of cortical areas are articulated more prominently in upper than deep layers (Charvet et al., 2015). For each region, the thickness for the upper layer compartments was estimated as the sum of the thickness of the following laminar compartments that existed in the matched von Economo and Koskinas areas: I, Ia, Ib, II, II+III, III, III(II), III(IV), IIIa, IIIa/b, IIIb, IIIb/c, IIIc. The cell density per mm^3^ was computed as the mean of these measurement for the upper laminar compartments. If for an area of the von Economo and Koskinas atlas information on thickness or cell density was available only for the dome or the wall of the area, this available information was used. Otherwise the mean values for the dome and wall measurements were taken into account. The same procedure was followed for the deep layer profiles by taking into account the laminar compartments V, V+VI, VI, VIa, VIb, Va, Vb. If more than one von Economo and Koskinas area was matched to a Desikan-Killiany region, the mean value of the structural traits of the von Economo and Koskinas areas was assigned to the Desikan-Killiany region (see Table 1).

The above procedure resulted in deep and upper layer estimates of the thickness and cell density per mm^3^ for each of the 34 regions of the Desikan-Killiany atlas. Thus, each region was characterized by four structural traits: deep and upper cortical thickness and deep and upper cortical cell density (for a schematic depiction see Figure 1 A, B).

In order to control the effect of the chosen specific parcellation on the results, we also performed the analysis for a connectome assembled on a high-resolution version of the Desikan-Killiany with 57 regions per hemisphere (see van den Heuvel et al., 2015a for details on the construction of this connectome).

### Relating structural traits to the existence of connections

The relation of the presence or absence of connections with the structural traits was examined by nominal logistic regression. The presence or absence of a connection between two regions served as the dependent variable. Hence, the dependent variable was binary, with 0 denoting absence and 1 denoting presence of a connection, irrespective of its strength. Since the connectivity matrix is undirected, only the upper triangle of the connectivity matrix was taken into account. The absolute difference of the four structural traits (deep and upper layer thickness and deep and upper layer cell density, respectively) of each pair of regions was used as a similarity measure. In addition, the physical distance between the pairs of regions was computed as the Euclidean distance between the mass centers of the regions. Physical distance is linked to the presence or absence of connections between brain areas, a reflection of the wiring cost principle (e.g., Kaiser and Hilgetag, 2006; Rubinov et al., 2015). In the case of connectomes constructed from diffusion imaging, physical distance is also known to affect the detection of connections between brains regions (e.g., Li et al., 2012). Hence, the evaluation of the role of physical distance for connections estimated from diffusion data, as in the current study, convolves a plausible neurobiological principle with biases of the imaging and tractography methods. Nonetheless, we included physical distance as an additional predictor. This approach allowed the examination of the relation of the structural traits and presence or absence of connections while controlling for the effect of distance.

As an alternative distance measure, we also used the fiber length distance. Fiber length distance is only available for existing connections. In order to approximate the “fiber length” of absent connections, support vector regression was used for modeling the relation of Euclidean distance between connected regions and the fiber length of the existing connections. The model was then applied to the Euclidean distances between unconnected regions for estimating the “fiber length” of absent connections.

The fit of the regression models was assessed by McFadden′s pseudo-R^2^. McFadden′s pseudo-R^2^ ranges from 0 to 1, with higher values indicating a better model (higher likelihood of the model when compared to the null model, that is, a model with only the intercept term). In order to assess the generalizability of the models, a prediction analysis was conducted. For assessing which predictor, or combination of predictors, led to maximum predictive power, nominal logistic regression models were build with all possible combinations of the independent variables (four structural traits plus physical distance, resulting in 31 models). The predictions were computed 100 times, each time using 80% of the available data to build the model and the remainder of the data serving as test data. The proportion of data used for training (varying from 80% to 90%) did not change the results. The quality of the predictions was assessed by computing receiver operating characteristic (ROC) curves and the corresponding areas under curve (AUC) for the original and null predictions (by shuffling the labels indicating which pair of areas were connected or not). The above analysis was conducted in the group average connectome for the left and right hemisphere separately.

We also sought to examine if any of the structural traits related to the strength of connections. Cytoarchitectonic similarity of cortical areas of the monkey and cat visual system is correlated with the strength of connections (i.e., number of axons) that link these areas (Hilgetag and Grant, 2010; Hilgetag et al., 2016). Diffusion imaging does not allow the reliable extraction of such quantitative information as compared to invasive tract-tracing methods. However, a moderate correlation of the number of tractography streamlines estimated from diffusion data and the underlying strength of connections as assessed by gold-standard invasive tract-tracing appears to exist (van den Heuvel et al., 2015b; Donahue et al., 2016). Hence, in the current context, we refer to connectivity strength as the number of diffusion tractography streamlines linking two regions. To uncover evidence for a relation of the strength of connections and structural traits, the strength was correlated with the similarity of deep and upper thickness and neuronal density difference of the interconnected regions. Spearman′s partial rank correlations were used and physical distance was also included as a variable. Thus, the computed correlations reflect the relation of connectivity strength with each structural trait (or physical distance) while controlling for physical distance and the remaining structural traits. The significance of the correlations was assessed via permutation statistics (i.e. shuffling the connectivity strengths). Null correlations were computed from 1000 permutations.

### Subject-wise versus group-wise analysis

A subject-wise analysis was carried out for assessing if the group average connectome analysis was representative and to ensure that the threshold involved in its construction did not bias the results. The same procedure as described above was carried out for each individual connectome. Since the structural traits were derived from the von Economo and Koskinas atlas, no subject-wise measurements could be obtained for the structural traits. Thus, the subject-wise analysis related each of the individual connectomes of the 215 subjects to the same structural traits as used in the group analysis.

To ensure that a specific choice of subject-wise thickness did not affect the results, we used the MRI-based subject-wise cortical thickness values. Cortical thickness is the only structural trait that can be directly measured with MRI and only for the cortical ribbon as a whole (at least with the current 3T MRI apparatus). Extraction of the MRI-based thickness values followed the procedure described in Scholtens et al., (2015). We used these MRI-based cortical thickness measurements of each subject and repeated the subject-wise analysis with the following predictors: Physical distance, upper and deep cell density differences, and MRI-based subject-wise cortical thickness differences (for the complete cortical mantle).

## Results

### Relation of structural traits to the existence of connections

The nominal logistic regression models using all factors simultaneously exhibited an overall good fit with a McFadden′s pseudo-R^2^ of 0.15 and 0.16 for the right and left hemisphere, respectively. Physical distance, as expected, was related to the presence or absence of connections in the group average connectome (beta=-5.35, -4.95, p<0.001, right and left hemisphere, respectively) as well as on a subject-wise level (215/215 (100%) subjects for both the left and right hemisphere; Figure 2). With respect to the structural traits, an interesting pattern was revealed: only the upper layer cell density difference reached statistical significance, for both hemispheres in the group average analysis (beta= -2.31, -2.07, p<0.01, right and left hemisphere respectively). No other structural trait reached significance (p>0.05) (Figure 2). For the subject-wise analysis, the upper layer cell density difference was significant (p<0.05) in 205/215 (95%) subjects for the right and 190/215 (88%) subjects for the left hemisphere (Figure 2). Other structural traits were significant only in a small number of subjects (fewer than 50/215 (23%) subjects) (Figure 2). Hence, upper cell density difference was consistently significantly related to connectivity both for the average connectome and subject-wise analysis, illustrating the close relation of this structural trait with the existence of connections between cortical areas.

**Figure 2.**
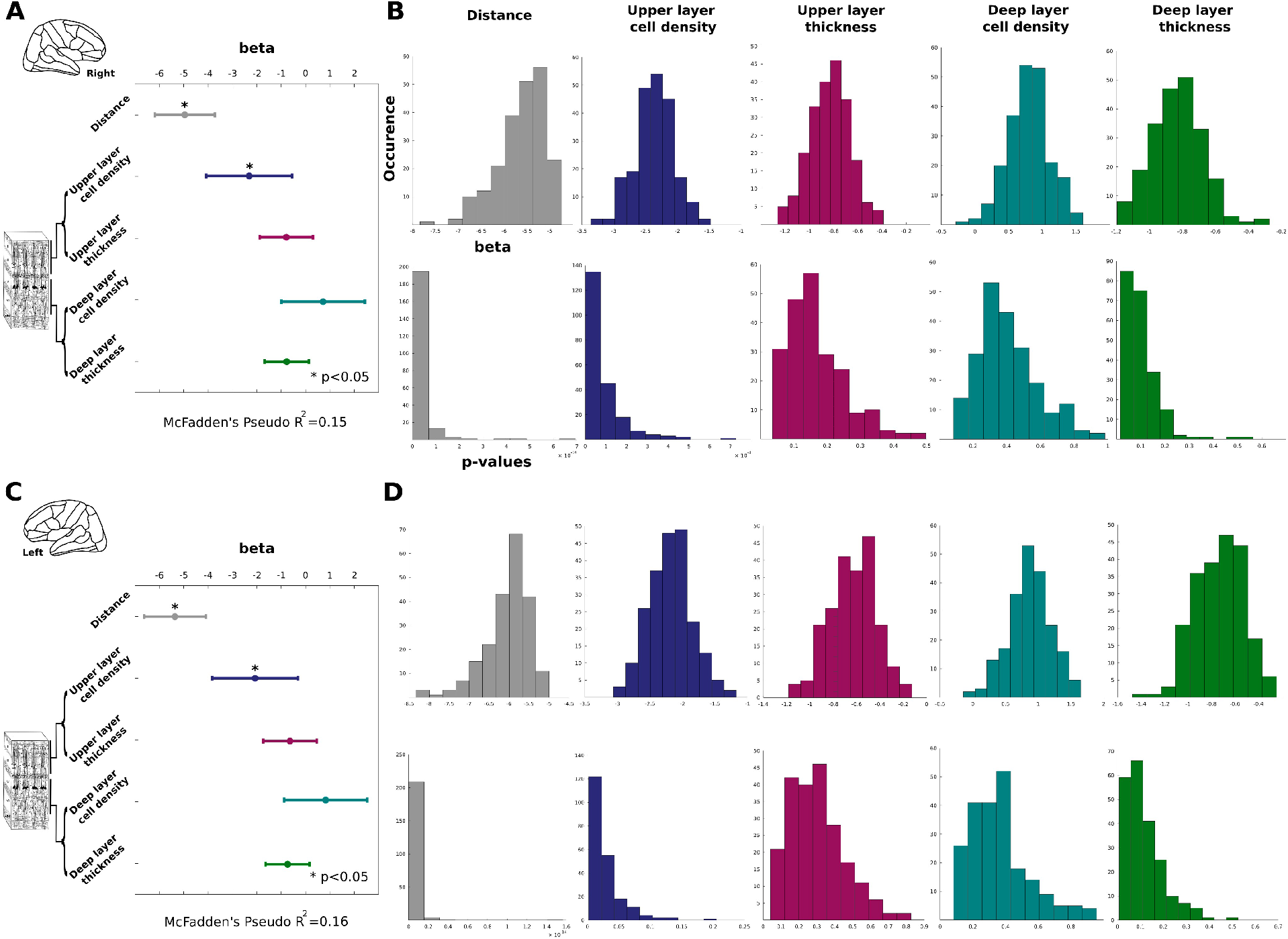
Logistic regression results on the group average connectome and the individual connectomes. **A.** For the right hemisphere, significant coefficients were obtained for distance (as expected) and upper (supragranular) cell density difference. **B .**Subject-wise analysis reveals the consistent association of distance and upper cell density difference with presence or absence of connectivity. Histograms depict the distribution of coefficients and respective p-values across 215 subjects. **C,D.** Same as A and B but for the left hemisphere analysis. Again distance and upper thickness difference consistently reaches significance.

Running the subject-wise analysis with the subject-wise MRI-based cortical thickness differences (in conjunction with the upper and deep cell density differences and distance predictors) showed the same picture. Distance and upper layer cell density difference were the only factors that reached significance (p<0.05). For distance, this was the case for 215/215 (100%) subjects for both the left and right hemisphere analysis. The upper layer cell density difference was found to be significant (p<0.05) in 207/215 (96%) subjects for the right hemisphere and 204/215 (94%) subjects for the left hemisphere. In contrast, the deep cell density difference and the MRI-based subject-wise cortical thickness difference was significant only in a small number of the subjects for both hemispheres (fewer than 40/215 (18%) subjects). All levels of significance for the subject-wise analysis were set to p<0.05 (uncorrected).

The prediction analysis also highlighted the upper layer cell density difference as the predictor leading to the highest performance in predicting presence and absence of connections. Specifically, upper layer cell density difference (in conjunction with physical distance, as expected) was the structural trait that was consistently part of the combination of features that led to the highest prediction performance for both the right and left hemisphere. For the right hemisphere, the highest prediction performance was achieved by the combination of distance, upper layer cell density difference, upper and deep thickness difference (AUC=0.76, p<0.01) followed by the combination of distance and upper layer cell density differences (AUC=0.75, p<0.01; Figure 3). For the left hemisphere, the highest prediction performance was achieved by the combination of distance, upper and deep layer cell density difference, followed by distance on its own (AUC=0.77, AUC=0.76, p<0.01 respectively). Moreover, when only combinations of structural predictors were taken into account, only upper layer cell density difference appeared consistently as the most predictive structural trait (in conjunction with deep thickness for the right hemisphere), leading to an AUC=0.63 and 0.60, p<0.01 for the right and left hemisphere, respectively (Figure 3). In addition, a sharp decrease in the AUC values was observed for combinations that did not include the upper layer cell density difference (Figure 3). Hence, upper layer cell density difference appeared as the only structural trait consistently part of the combinations that led to better predictions of the existence of connections for both the right and left hemisphere.

**Figure 3.**
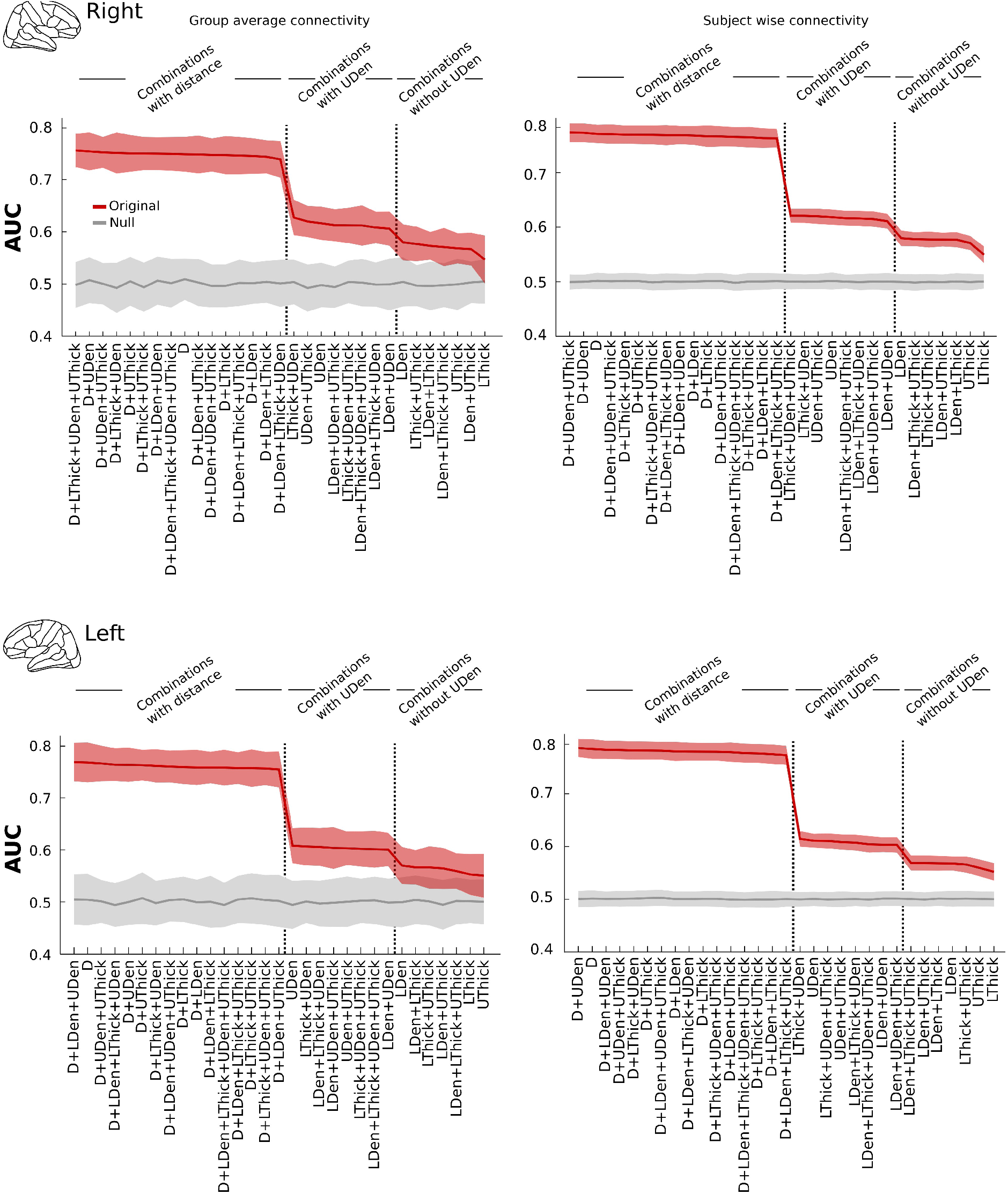
Prediction analysis with all possible combinations of predictors. Red color indicates the original AUC values with thick lines indicating the mean values and the shaded region the standard deviations across 100 cross-validations. Grey color denotes null values derived from predictions with shuffled presence/absence connectivity labels. upper cell density difference is the only structural trait that conjointly with distance leads to the highest AUC values. Moreover, when only combinations of structural traits are considered, upper cell density difference is again consistently the structural trait that leads to the highest AUC values. In addition, note that a sharp decrease in the AUC values is observed for models not including the upper cell density difference similarity of regions. The shaded areas for the group average analysis represent AUC standard deviation across 100 cross-validation steps, whereas for the subject-wise analysis they represent AUC standard deviation across the analysis performed in 215 subjects. All the aforementioned AUC values are statistically significant (p<0.01). Abbreviations: D: Distance, DDen: deep (infragranular) cell density difference, DThick: deep (infragranular) thickness difference, UDen: upper (supragranular) cell density difference, UThick: upper (supragranular) thickness difference.

The prediction analysis conducted on a subject-wise basis revealed the same picture as the group average analysis; that is, the upper layer cell density difference was the one structural trait that was consistently part of the combination of features leading to a high prediction performance. Specifically, for the right hemisphere, the upper layer cell density difference in conjunction with upper thickness and physical distance led to an AUC=0.77, p<0.01. For the left hemisphere, upper layer cell density difference in conjunction with distance led to an AUC=0.79, p<0.01. Furthermore, when only structural traits were used as features, upper cell density appeared again as the feature consistently part of feature combinations with the greatest predictive power (in conjunction with deep and upper layer thickness for the right hemisphere AUC=0.62, p<0.01, and in conjunction with deep layer thickness for the left hemisphere AUC=0.61, p <0.01).

Using fiber length as a measure of wiring distance led to qualitatively the same results. Specifically, the logistic regression for the left hemisphere resulted in a good fit (McFadden′s pseudo-R^2^ =0.18) significant values only for physical distance (beta=-5.70) and upper layer cell density similarities (beta=-1.79) (all p<0.01). For the right hemisphere the logistic regression resulted in a good fit (McFadden′s pseudo-R^2^ =0.16) significant values only for physical distance (beta=-5.12) and upper layer cell density similarities (beta=-2.26) (all p<0.01). The prediction analysis for the left hemisphere revealed again that among the structural traits, upper layer cell density similarity was consistently part of the combinations leading to the highest AUC values (higher AUC=0.78 for physical distance, followed by AUC=0.77 in conjunction with upper layer cell density). The same held true for the right hemisphere (AUC=0.76 for the combination of physical distance and upper layer cell density difference).

Moreover, the results for the high resolution Desikan-Killiany parcellation were qualitatively the same. Specifically, the logistic regression for the left hemisphere resulted in a good fit (McFadden′s pseudo-R^2^ =0.22), significant values only for physical distance (beta=-7.39) and upper layer cell density similarities (beta=-1.80) (all p<0.01). For the right hemisphere, the logistic regression resulted in a good fit (McFadden′s pseudo-R^2^ =0.20), significant values only for physical distance (beta=-7.48) and upper layer cell density similarities (beta=-1.25) (all p<0.01). The prediction analysis for the left hemisphere revealed again that, among the structural traits, upper layer cell density similarity was consistently part of the combinations leading to the highest AUC values (higher AUC=0.82 for physical distance with upper cell density and deep cell thickness). The same held true for the right hemisphere (AUC=0.81 for the combination of physical distance, upper and deep layer cell density difference).

Apart from physical distance, which as expected exhibited a strong negative correlation with the number of the streamlines (i.e., connection strength) between two regions, a modest correlation was observed only with respect to upper layer cell density similarity (Figure S1). This finding also held true when fiber length was used as wiring distance (left hemisphere rho=-0.47, -0.30, right hemisphere rho=-0.54, -0.26, for physical distance and upper layer cell density similarity, all p<0.001). This was also the case for the high resolution Desikan-Killiany parcellation (left hemisphere rho=-0.60, -0.15, right hemisphere rho=-0.60, -0.20, for distance and upper cell density similarity, all p<0.01).

Taken together, the above results indicate that the upper layer cell density difference is consistently related to the existence of connections between cortical regions. Importantly, this finding held true for both the left and right hemisphere analysis, for the group-wise and subject-wise analysis, the use of fiber length as a proxy for wiring cost, as well as for a different parcellation scheme. By contrast, the other structural traits did not appear to be consistently and closely linked to the existence of connections. In addition, a modest negative correlation was observed between the upper layer cell density similarity and the number of streamlines running between two regions, but not for other structural traits (Figure S1).

## Discussion

### Upper layer cell density similarity of cortical regions is related to the existence of connections

One major goal of human brain research is the detailed mapping and understanding of the macroscale wiring of the human brain. This scientific endeavor has produced numerous insights into the topological properties of the human cortical connectome, such as the existence, functional significance and vulnerability of hubs. However, it is still unclear what principles govern this wiring diagram at a basic level; for instance, what mechanisms systematically explain the existence of a connection between some areas, but not others. Only recently have studies begun to address this fundamental question and tried to unveil wiring principles beyond the well documented aspect of wiring cost (Costa et al., 2007; Vértes et al., 2012; Samu et al., 2014; Betzel et al., 2016). Our previous work on the wiring of the mammalian cortex (in mouse, cat and macaque monkey) demonstrated that cytoarchitectonic similarity is predictive of the presence or absence of cortico-cortical connections (Barbas et 2005; Beul et al., 2015a; Goulas et al., in press; Beul et al., 2015b; Hilgetag et al., 2016). Applying this approach to the connectional architecture of the human cortex, we here demonstrate that cytoarchitecture is also closely linked to the existence of connections in the human brain. More specifically, it is the cell density similarity of the upper laminar compartments of cortical areas that is essential for predicting connectional features.

The neuroanatomists of the classical era considered brain structure and function as a whole, and consistently approached neuromorphology in a histophysiological perspective, and not as a mere descriptive parcellation of cellular populations (Ramón y Cajal, 1900–1906; Jakob, 1939–1941).Pioneers of cortical histoarchitecture, such as Ramón y Cajal, Brodmann, von Economo and Koskinas, and the Vogts, worked on the premise that morphological diversity reflects functional specifications: “As a general principle, each physiological function presupposes a corresponding anatomical basis. From the precise knowledge of the structure of the cerebral cortex we may expect to shed light on issues of the utmost importance, such as the relationship between mental attributes and brain structure” (Koskinas, 1931). Thus, cortical cytoarchitecture and myeloarchitecture are inextricable from cortical connections. Areas that are primarily sensory exhibit dense layers II and IV rich in small-diameter granule cells that are local interneurons. On the other hand, cytoarchitectonic areas with motor specifications contain large neurons in layers III and V, as these neurons need large somata to sustain their long projecting axons. The relation of soma size and axon diameter has been explicitly demonstrated for the contralateral connections of the macaque prefrontal and motor cortex (Tomasi et al., 2012).

Why does the similarity of upper, but not deep layer, cell density of cortical areas carry information on connectivity? The formation of the cortical layers proceeds in an “inside-out” fashion with neuron migration finishing in less eulaminated areas (e.g., cingulate cortex) earlier than in more eulaminated areas (e.g., primary visual cortex) (Rakic, 2002). Therefore, the characteristic upper cortical layers of eulaminate areas, including a well-developed granular layer 4, are a result of late developmental periods, while deep cortical layers are formed in all areas early in development, and, thus, are less well distinguished from each other.

It is plausible that the mechanism giving rise to the relation between cytoarchitecture and connectivity is related to the co-occurrence of the processes of neurogenesis and neuronal migration as well as the formation of axonal projections during distinct time windows of the development of the different cortical areas (Dombrowski et al., 2001; Barbas et al., 2015). This scenario suggests that connections are more likely to be formed between cortical areas that are populated during the same or overlapping developmental time windows, since they contain more neuronal ‘partners’ available for establishing connections. As a consequence, upper cell density differences represent a proxy of the similarity of developmental time windows of cortical areas, and consequently for the likelihood of establishing connections between them. In summary, while previous animal studies demonstrated that upper cell densities characteristically reflect cytoarchitectonic gradients across the cortical sheet, our current results suggest an additional prominent role of upper cell density in the macroscale wiring of the human cortex.

The lack of a significant and systematic relation of cortical thickness with the existence of connections is in line with previous work on the macaque cortex (Beul et al., 2015b; Hilgetag et al., 2016). Despite the fact that thickness and cytoarchitecture of cortical areas tend to covary across the cortical sheet (e.g., von Economo, 1927), the thickness similarity of cortical areas does not carry additional information on the existence and strength of connections and might not be a good proxy for neurodevelopmental events that are important for the establishment and maintenance of cortico-cortical connections.

### Wiring principles and neurobiological mechanisms

The current results demonstrate that the cytoarchitectonic similarity of upper cortical layers is closely linked to the existence of cortical connections, and appears to underlie a basic wiring principle of the human connectome. However, the findings do not directly unveil the neurobiological mechanisms that give rise to the observed relationship. While we sketched out a plausible neurodevelopmental mechanism, further empirical and modeling studies are necessary to understand the details of the developmental relationships. The value of identifying wiring principles lies in establishing a quantitative ground, providing the starting point for the development of hypotheses on neurodevelopmental mechanisms. This direction is different from approaches that aim merely at maximizing the prediction performance of features, for instance, for the existence of connections (e.g., Hinne et al., 2016). Hence, a meaningful wiring principle should not only possess a high correlation with connectional features, but also the potential of a neurobiological interpretation.

### Relation to orderly spatial patterns of cortico-cortical connectivity

A prominent feature of cortical architecture is the presence of orderly spatial patterns of area-to-area connections (in which neighboring source areas connect primarily to neighboring target areas), that is, connection topography. For instance, patterns of connections among areas of the macaque monkey visual system obey the spatial arrangement dictated by the retinotopy of the visual areas (e.g., Kaas, 2002). Orderly spatial patterns of connections also exist between non-primary sensory areas, such as the prefrontal and temporal cortex or the frontal and parietal cortex of the macaque monkey (see for instance figures 17 B and 19 B in Pandya and Yeterian, 1990). Orderly spatial patterns of connections are also observed in the human brain. For instance, functional connectivity between the ventral frontal cortex and parietal cortex, as estimated by resting state fMRI, exhibit a spatially ordered pattern centered around the central sulcus (Margulies and Petrides, 2013).

Why do such orderly spatial patterns characterize certain cortico-cortical connections? The answer might lie in the fact that the cytoarchitecture of cortical areas, that is the degree of their eulamination, is a spatially ordered gradient that reflects the so-called “Gradationsprinzip” (gradation principle, Vogt and Vogt, 1919; Sanides, 1962). Our findings demonstrate that in the human brain a “similar prefers similar” cytoarchitecture-based principle is closely related to cortico-cortical connectivity. This wiring principle in conjunction with the gradation principle of cytoarchitectonic organization offers a fundamental framework with the potential for systematically explaining the spatially ordered patterns of connectivity that are observed in various studies of human cortical connectivity. Based on this framework, if two areas in different parts of the cerebral sheet have similar cytoarchitecture, it is likely that they will be connected. The gradation principle dictates that their neighbouring areas will also have a similar cytoarchitecture and therefore will also likely be connected. Thus, the combination of the similarity and gradation principles results in a spatially ordered connectivity pattern. In other words, distributed long-range connections, as well as spatial motifs of resting-state networks (Power et al., 2011), might possess orderly spatial patterns due to the spatially graded cytoarchitecture of the cortical sheet and the fact that cytoarchitectonic similarity constitutes a wiring principle. Therefore, the current results can offer a systematic explanation for aspects of the “topographic human connectome” (Jbabdi et al., 2013).

### Limitations and future directions

The cortico-cortical connectome used in the present analysis was estimated from high quality diffusion imaging data of the Human Connectome Project. Despite the fact that this dataset is based on state-of-the art acquisition and prepossessing strategies, connectomes assembled from diffusion imaging are very likely to include false positive and negative information (e.g., Thomas et al., 2014). Long-range connections suffer particularly strongly from such deficits (Li et al., 2012). Therefore, it is likely that connections linking areas with similar cytoarchitecture in different cortical lobes are missing from studies based on current *in vivo* techniques, potentially leading to an under-appreciation of the role of cytoarchitecture in the human connectome. In other words, the predictive power of cytoarchitecture might be even higher than demonstrated by our analyses. Such methodological limitations might also contribute to the observation that cortical thickness appeared as a negligible structural trait. However, our prior work with connectivity assessed by gold-standard invasive tract-tracing techniques in the macaque monkey (Beul et al., 2015b, Hilgetag et al., 2016) cortex argues against such an interpretation. These analyses of data for the non-human primate cortex delivered very similar findings as made here for the human cortex, showing a fundamental relation of cytoarchitecture to connectivity and a negligible role of cortical thickness.

Another limitation of the current approach is the matching of the regions of the Desikan-Killiany atlas to the cortical areas of von Economo and Koskinas (1925) and the extrapolation of the measurements provided in von Economo and Koskinas (1925) to these regions. This convolution represents an approximation, since the von Economo and Koskinas areas for which quantitative information is available do not map to the 34 Desikan-Killiany regions in a one-to-one fashion (e.g., areas partially overlap and do not exactly match the Desikan-Killiany regions). Recent efforts such as the BigBrain project (Amunts et al., 2013) might eventually provide the opportunity to extract detailed quantitative cytoarchitectonic information directly from a modern stereotaxic atlas that can be spatially co-registered to diffusion imaging data.

### Conclusions

We demonstrated that the cytoarchitectonic similarity of cortical areas in the human brain, specifically the similarity of cell density of upper cortical layers, is tightly linked to the presence or absence of cortico-cortical connections. In conjunction with our previous studies on the mouse, cat and macaque cortex, the current findings highlight cytoarchitectonic similarity as an overarching principle underlying the wiring of the mammalian cortex. This principle forms the foundation for hypotheses about neurodevelopmental mechanisms for the establishment and maintenance of connections, as well as an explanation for the spatially orderly connectivity patterns observed in the human cerebral cortex.

## Acknowledgements

Supported by a Humboldt Research Fellowship from the Alexander von Humboldt Foundation to AG, and grants of the DFG (SFB 936/ A1, Z3; TRR 169/ A2) to CCH. Human neuroimaging data were kindly provided by the Washington University, University of Minnesota, and Oxford University (WU-Minn HCP consortium; Principal Investigators: David Van Essen and Kamil Ugurbil; Grant 1U54MH091657) funded by the 16 National Institutes of Health (NIH) institutes and centers that support the NIH Blueprint for Neuroscience Research and by the McDonnell Center for Systems Neuroscience at Washington University.

---

**Figure S1.**
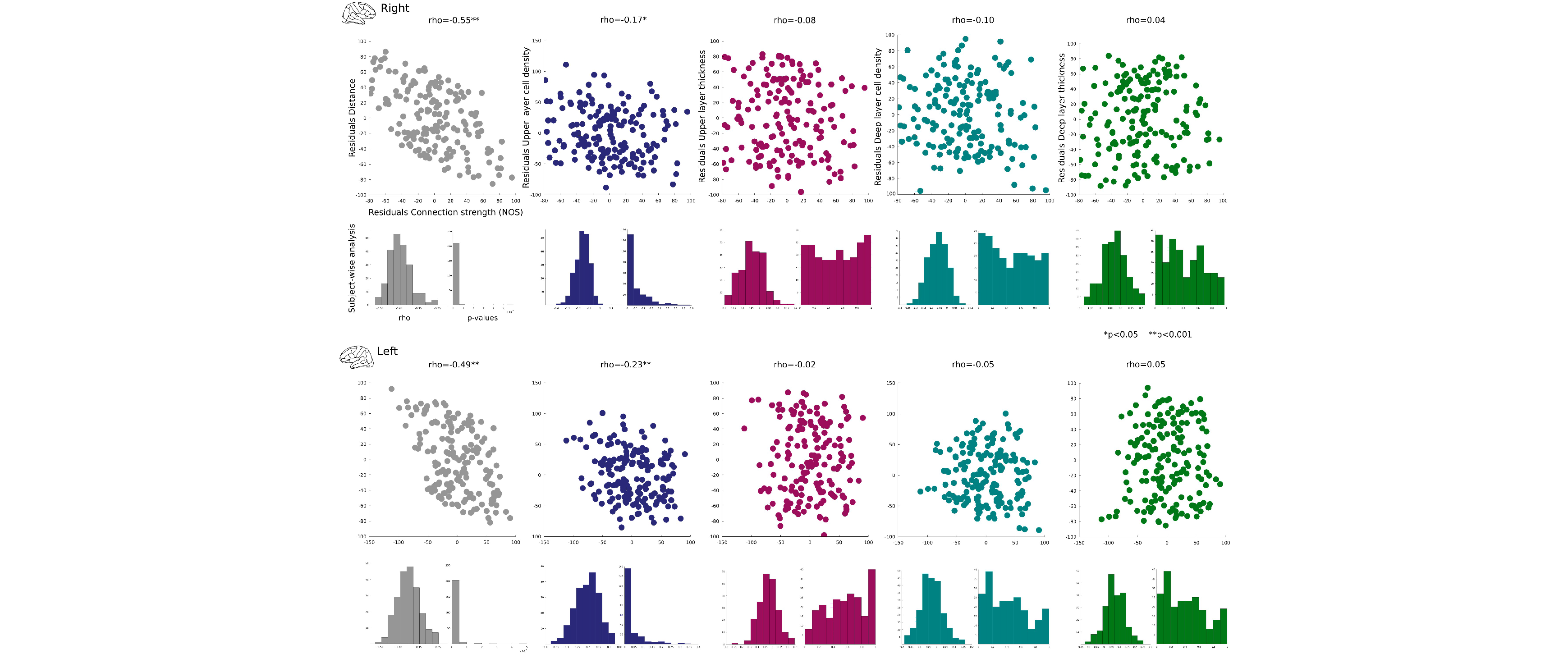
Correlations of connectivity strength and structural traits. Correlations are partial Spearman′s rank correlations. Hence, the scatterplots depict the relation between the residuals of connectivity strength and each structural trait after regressing out the rest of the structural traits. The only significant correlations of connectivity strength that were observed was, as expected, with distance and with upper (supragranular) neuronal density difference. Significance of correlations was established with 1000 permutations.

## References

Amunts K, Lepage C, Borgeat L, Mohlberg H, Dickscheid T, Rousseau ME, Bludau S, Bazin PL, Lewis LB, Shah NJ, Lippert T, Zilles K, Evans AC (2013). BigBrain: An ultra high-resolution 3D human brain model. Science 340:1472-1475.

Barbas H, Hilgetag CC, Saha S, Dermon CR, Suski JL (2005). Parallel organization of contralateral and ipsilateral prefrontal cortical projections in the rhesus monkey. BMC Neurosci 6:32.

Barbas H (2015). General cortical and special prefrontal connections: principles from structure to function. Annu Rev Neurosci 38:269–289.

Betzel RF, Avena-Koenigsberger A, Goñi J, He Y, de Reus MA, Griffa A, Vértes PE, Mišic B, Thiran J-P, Hagmann P, van den Heuvel M, Zuo X-N, Bullmore ET, Sporns O (2016). Generative models of the human connectome. NeuroImage 124:1054–1064.

Beul SF, Grant S, Hilgetag CC (2015a). A predictive model of the cat cortical connectome based on cytoarchitecture and distance. Brain Struct Funct 220:3167–3184.

Beul SF, Barbas H, Hilgetag CC (2015b). A predictive structural model of the primate connectome. arXiv:1511.07222.

Bullmore E, Sporns O (2009). Complex brain networks: graph theoretical analysis of structural and functional systems. Nat Rev Neurosci 10:186–198.

Brodmann K (1909). Vergleichende Localizationslehre der Grosshirnrinde in ihren Prinzipien dargestellt auf Grund des Zellenbaues. Leipzig: Barth.

Campbell AW (1905). Histological studies on the localisation of cerebral function. Cambridge: University Press.

Costa LF, Kaiser M, Hilgetag CC (2007). Predicting the connectivity of primate cortical networks from topological and spatial node properties. BMC Syst Biol. 1:16.

Crossley N, Mechelli A, Scott J, Carletti F, Fox PT, McGuire P, Bullmore ET (2014). The hubs of the human connectome are generally implicated in the anatomy of brain disorders. Brain 137:2382–2395.

Charvet CJ, Cahalane DJ, Finlay BL (2015). Systematic, cross-cortex variation in neuron numbers in rodents and primates. Cereb Cortex 25:147–160.

Desikan RS, Ségonne F, Fischl B, Quinn BT, Dickerson BC, Blacker D, Buckner RL, Dale AM, Maguire RP, Hyman BT, Albert MS, Killiany RJ (2006). An automated labeling system for subdividing the human cerebral cortex on MRI scans into gyral based regions of interest. NeuroImage 31:968–980.

Dombrowski SM, Hilgetag CC, Barbas H (2001). Quantitative architecture distinguishes prefrontal cortical systems in the rhesus monkey. Cereb Cortex. 11:975–988.

Donahue CJ, Sotiropoulos SN, Jbabdi S, Hernandez-Fernandez M, Behrens TE, Dyrby TB, Coalson T, Kennedy H, Knoblauch K, Van Essen DC, Glasser MF (2016). Using diffusion tractography to predict cortical connection strength and distance: a quantitative comparison with tracers in the monkey. J Neurosci 36:6758-70.

de Reus MA, van den Heuvel MP (2013). Estimating false positives and negatives in brain networks. NeuroImage 70:402-9.

Eickhoff S, Stephan KE, Mohlberg H, Grefkes C, Fink GR, Amunts K, Zilles K (2005). A new SPM toolbox for combining probabilistic cytoarchitectonic maps and functional imaging data. NeuroImage 25:1325-1335.

Fischl B, Dale AM (2000). Measuring the thickness of the human cerebral cortex from magnetic resonance images. Proc Natl Acad Sci USA 97:11050-11055.

Fischl B, Van Der Kouwe A, Destrieux C, Halgren E, Ségonne F, Salat DH, Busa E, Seidman LJ, Goldstein J, Kennedy D, Caviness V, Makris N, Rosen B, Dale AM (2004). Automatically parneuronalating the human cerebral cortex. Cereb Cortex 14:11:22.

Goulas A, Uylings HBM, Hilgetag CC (in press) Principles of ipsilateral and contralateral cortico-cortical connectivity in the mouse. Brain Struct Funct.

Hagmann P, Cammoun L, Gigandet X, Meuli R, Honey CJ, Wedeen VJ, Sporns O (2008). Mapping the structural core of human cerebral cortex. PLoS Biol 6:e159.

Hilgetag CC, Grant S (2010). Cytoarchitectural differences are a key determinant of laminar projection origins in the visual cortex. NeuroImage 51:1006–1017.

Hinne M, Meijers A, Bakker R, Tiesinga PHE, Mørup M, van Gerven MAJ (2016). The missing link: predicting connectomes from noisy and partially observed tract tracing data. bioRxiv doi: http://dx.doi.org/10.1101/063867.

Jakob C (1939–1941). El Cerebro Humano: Atlas I–III. Buenos Aires: Aniceto López.

Jbabdi S, Sotiropoulos S, Behrens T (2013). The topographic connectome. Curr Opin Neurobiol 23:207–215.

Kaas JH (2002). Cortical areas and patterns of cortico-cortical connections. In: Schüz A, Miller R (eds) Cortical areas: Unity and diversity. Taylor and Francis, p.179-195.

Kaiser M, Hilgetag CC (2006). Nonoptimal component placement, but short processing paths, due to long-distance projections in neural systems. PLoS Comput Biol 2:e95.

Koskinas GN (1931). Scientific works Published in German. Athens: Pyrsus Publishers.

Li L, Rilling JK, Preuss TM, Glasser MF, Hu X (2012). The effects of connection reconstruction method on the interregional connectivity of brain networks via diffusion tractography. Hum Brain Mapp 33:1894–1913.

Margulies DS, Petrides M (2013). Distinct parietal and temporal connectivity profiles of ventrolateral frontal areas involved in language production. J Neurosci 33:16846–16852.

Medalla M, Barbas H (2006). Diversity of laminar connections linking periarcuate and lateral intraparietal areas depends on cortical structure. Eur J Neurosci 23:161–179.

Meunier D, Lambiotte R, Fornito A, Ersche KD, Bullmore ET (2009). Hierarchical modularity in human brain functional networks. Front Neuroinform 3:37.

Pandya DN, Yeterian EH (1985). Architecture and connections of cortical association areas. In: Peters A, Jones EG. Association and Auditory Cortices. Springer Science Business Media New York, p.3-61.

Pandya DN, Seltzer B, Barbas H (1988). Input-output organiztion of the primate cerebral cortex. In: Steklis H, Erwin J (eds) Comparative Primate Biology, Vol.4: Neurosciences. Alan R. Liss, New York, p. 39–80.

Pandya DN, Yeterian EH (1990). Prefrontal cortex in relation to other cortical areas in rhesus monkey: architecture and connections. In: Uylings HBM, van Eden CG, de Bruin JPC, Feenstra MGP, Pennartz CMA, editors. The prefrontal cortex: its structure, function and pathology. Progress in Brain Research. Vol. 85. Amsterdam: Elsevier. p. 63–94.

Pandya DN, Petrides M, Seltzer B, Cipolloni BP (2015). Cerebral cortex: Architecture, connections, and the dual origin concept. New York: Oxford University Press.

Power JD, Cohen AL, Nelson SM, Wig GS, Barnes KA, Church J, Vogel AC, Laumann TO, Miezin FM, Schlaggar BL, Petersen SE (2011). Functional network organization of the human brain. Neuron 72:665–678.

Rakic P (2002). Neurogenesis in adult primate neocortex: an evaluation of the evidence. Nat Rev Neurosci. 3:65-71.

Ramón y Cajal S (1900–1906). Studien über die Hirnrinde des Menschen, 1.–5. Heft (transl. by J. Bresler). Leipzig: Barth.

Rubinov M, Ypma RJF, Watson C, Bullmore ET (2015). Wiring cost and topological participation of the mouse brain connectome. Proc Natl Acad Sci 112:10032-10037.

Samu D, Seth AK, Nowotny T (2014). Influence of wiring cost on the large-scale architecture of human cortical connectivity. PLoS Comput Biol 10:e100355.

Sanides F (1962). Die Architectonik des menschlichen Stirnhirns. Berlin-Heidelberg: Springer-Verlag.

Senden M, Deco G, de Reus M, Goebel R, van den Heuvel MP (2014). Rich club organization supports a diverse set of functional network configurations. NeuroImage 96:174–182.

Scholtens LH, Schmidt R, de Reus MA, van den Heuvel MP (2014). Linking macroscale graph analytical organization to microscale neuroarchitectonics in the macaque connectome. J Neurosci 34:12192–12205.

Scholtens LH, de Reus M, van den Heuvel MP (2015). Linking contemporary high resolution magnetic resonance imaging to the von Economo legacy: A study on the comparison of MRI cortical thickness and histological measurements of cortical structure. Hum Brain Mapp 36:3038–3046.

Thomas C, Ye FQ, Irfanoglu MO, Modi P, Saleem KS, Leopold D, Pierpaoli C (2014). Anatomical accuracy of brain connections derived from diffusion MRI tractography is inherently limited. Proc Natl Acad Sci USA 111:16574-16579.

Tomasi S, Caminiti R, Innocenti GM (2012). Areal differences in diameter and length of corticofugal projections. Cereb Cortex 22:1463–1472.

van den Heuvel MP, Sporns O (2013). Network hubs in the human brain. Trends Cogn Sci 17:683–696.

van den Heuvel MP, Scholtens LH, Feldman Barrett L, Hilgetag CC, de Reus M (2015a). Bridging cytoarchitectonics and connectomics in human cerebral cortex. J Neurosci 35:13943–13948.

van den Heuvel MP, de Reus MA, Feldman Barrett L, Scholtens LH, Coopmans FM, Schmidt R, Preuss TM, Rilling JK, Li L (2015b). Comparison of diffusion tractography and tract-tracing measures of connectivity strength in rhesus macaque connectome. Hum Brain Mapp 36:3064-3075.

Van Essen DC, Smith SM, Barch DM, Behrens TEJ, Yacoub E, Ugurbil K (2013). The WU-Minn Human Connectome Project: an overview. NeuroImage 80:62–79.

Vértes PE, Alexander-Bloch AF, Gogtay N, Giedd JN, Rapoport JL, Bullmore ET (2012). Simple models of human brain functional networks. Proc Natl Acad Sci U S A 109:5868–5873.

Vogt C, Vogt O (1919). Ergebnisse unserer hirnforschung. 1-4. Mitteilung. J. Psychol. Neurol.25:279–461.

von Economo C (1927). Zellaufbau der Grosshirnrinde des Menschen. Berlin: Springer.

von Economo C, Koskinas GN (1925). Die Cytoarchitektonik der Hirnrinde des erwachsenen Menschen. Wien:Springer.

Zamora-López G, Zhou C, Kurths J (2010). Cortical hubs form a module for multisensory integration on top of the hierarchy of cortical networks. Front Neuroinform 4:1.

